# Plant functional types and tissue stoichiometry explain nutrient transfer in common arbuscular mycorrhizal networks of temperate grasslands

**DOI:** 10.1101/2022.10.05.511035

**Authors:** Hilary Rose Dawson, Katherine L. Shek, Toby M. Maxwell, Paul B. Reed, Barbara Bomfim, Scott D. Bridgham, Brendan J. M. Bohannan, Lucas C.R. Silva

**Affiliations:** Institute of Ecology and Evolution, University of Oregon, Eugene, OR 97403 USA; Department of Biology, University of Oregon, Eugene, OR 97403 USA; Research School of Biology, Australian National University, Canberra, ACT 2601 Australia; Department of Natural Resources and the Environment, University of New Hampshire, Durham NH 03824; Department of Biological Sciences, Boise State University, Boise, Idaho, 83725 USA; Institute for Applied Ecology, Corvallis, OR 97333 USA; Climate and Ecosystem Sciences Division, Lawrence Berkeley National Laboratory, Berkeley, CA 94720 USA; Environmental Studies Program, University of Oregon, Eugene, OR 97403 USA

**Author notes:** **Corresponding authors**: H.R.D.; L.C.R.S.

**Keywords:** arbuscular mycorrhizal fungi, carbon, common mycorrhizal networks, ecological stoichiometry, grasslands, nitrogen, plant functional type

## Abstract

Plants and mycorrhizal fungi form mutualistic relationships that affect how resources flow between organisms and within ecosystems. Common mycorrhizal networks (CMNs) could facilitate preferential transfer of carbon and limiting nutrients, but this remains difficult to predict. Do CMNs favor fungal resource acquisition at the expense of plant resource demands (a fungi-centric view), or are they passive channels through which plants regulate resource fluxes (a plant-centric view)? We used stable isotope tracers (^13^CO_2_ and ^15^NH_3_), plant traits, and mycorrhizal DNA to quantify above- and belowground carbon and nitrogen transfer between 18 plant species along a 520-km latitudinal gradient in the Pacific Northwest, USA. Plant functional type and tissue stoichiometry were the most important predictors of interspecific resource transfer. Of “donor” plants, 98% were ^13^C-enriched, but we detected transfer in only 2% of “receiver” plants. However, all donors were ^15^N-enriched and we detected transfer in 81% of receivers. Nitrogen was preferentially transferred to annuals (0.26 ± 0.50 mg N per g leaf mass) compared to perennials (0.13 ± 0.30 mg N per g leaf mass). This corresponded with tissue stoichiometry differences. Our findings suggest that plants and fungi that are located closer together in space and with stronger demand for resources over time are more likely to receive larger amounts of those limiting resources.

## Introduction

Plant-mycorrhizal associations are thought to have emerged as rudimentary root systems over 400 million years ago, facilitating the expansion of terrestrial life that followed (Kenrick & Strullu-Derrien, 2014). The transformative power of early fungal symbioses is still evident today in all major plant lineages, from bryophytes to angiosperms (Heijden et al., 2015). Over 85% of all contemporary flowering plant species form symbioses with fungi, with arbuscular mycorrhizal (AM) associations being the most common (Brundrett, 2009). The relationships between plants and AM fungi dominates both managed and unmanaged landscapes and are estimated to be responsible for up to 80% of global primary productivity (Heijden et al., 2015). Fungi can form symbioses with more than one individual plant, particularly AM fungi which have low host specificity (Selosse et al., 2006). By extension, it has been widely hypothesized that the multi-plant-fungal relationships form “common mycorrhizal networks” (CMNs) which facilitate carbon and nutrient transfer between organisms, beyond the immediate plant-fungus mutualism formed by individuals. Although CMNs are traditionally defined by strict criteria that are difficult to test experimentally (Karst et al., 2008), the concept is ecologically relevant and useful in designing new experiments that may bring insight into CMNs structure and function. Here, we use ‘CMN’ under the proposed new definition of Rillig et al (2024) “where at least one mycorrhizal fungal genet interacts (connecting and colonizing or growing in close proximity) with the roots of a minimum of two plants of the same or different species.” Although Rillig et al. (2024) claim it is unlikely that carbon and nutrient exchanges will occur without hyphal continuity, we disagree that hyphal continuity is necessary for these exchanges. Plants leak exudates into the soil that can be picked up by fungal hyphae near the root surface (Vives-Peris et al., 2020). The same fungi can be in symbiosis with multiple plants. Although this interaction is not what is classically considered a CMN (what Rillig et al. have renamed as a CMN with hyphal continuity, or CMN-HC), it remains in essence fungal transport of carbon and nutrients derived from the plants involved (a CMN by Rillig et al.’s revised definition). Both CMN-HCs and CMNs could have ecological significance. The CNM relationships are often simplified to the mutual exchange of carbon-based photosynthates for soil nutrients that are more readily bioavailable to fungi (Smith & Read, 2008), but exist along a continuum from parasitic to mutualistic (Johnson et al., 1997; Karst et al., 2008; Luo et al., 2023).

The literature holds myriad and often complimentary, but sometimes contradictory, hypotheses that could explain the CMN mutualism as a key structural and functional component of ecosystems. For example, the “economics” hypothesis (Kiers et al., 2011) proposes that plants and fungi engage in “trades” of nutrients mined by fungi in exchange for plant photosynthates (Averill et al., 2019; Fellbaum et al., 2014; Werner & Dubbert, 2016). In the economics hypothesis, the terms of trade between plant and fungi are mediated by supply and demand for limiting resources, which could create a dynamic market emerging from interactions between environmental, biochemical, and biophysical variables. The “Wood Wide Web” hypothesis emerged from the analysis of isotopically labeled carbon transferred between plants, presumably through fungal mycorrhizae. Simard et al. (1997) hypothesized that plants that allocate carbon to sustain common fungal symbionts also benefit from shared nutrients, while plants associating with mycorrhizal fungi outside that network cannot. Complementing the analogy, the “kinship” hypothesis proposes that plants of the same species preferentially receive more resources in CMNs (Pickles et al., 2017; Tedersoo et al., 2020).

The past two decades have seen extensive but inconclusive research on these hypotheses and how they relate to empirical measurements of CMN structure and function. On the one hand, economic analogies suggest that the reciprocally regulated exchange of resources between plants and fungi in CMNs should favor the most beneficial cooperative partnerships (Fellbaum et al., 2014; Kiers et al., 2011). On the other hand, reciprocal transfer is only found in a subset of symbionts under specific conditions, while increased competition in CMNs is a more common observation (Walder & Van Der Heijden, 2015; Weremijewicz et al., 2016). At the core of this controversy is whether CMNs actively support fungal resource acquisition at the expense of plant resource demands (i.e., a fungi-centric view) or function as passive channels through which plants regulate resource fluxes (i.e., a plant-centric view). If plant-centric, we expect to find that the structure and functioning of CMNs give rise to consistent spatiotemporal patterns of resource allocation similar to those predicted by the kinship hypothesis. If fungi-centric, we expect to find that spatiotemporal patterns of resource allocation reflect the composition and functioning of the fungal community regardless of the connecting plant nodes in CMNs. Other perspectives emphasize that CMNs are experimentally under-documented and that this is an area that warrants further research (Henriksson et al., 2023; Karst et al., 2023; Rillig et al., 2024; Robinson et al., 2024). Given that data exist to support multiple, sometimes opposing views (Figueiredo et al., 2021; Silva & Lambers, 2021); we posit that CMNs are neither plant- nor fungi-centric.

In this study, we ask if interactions among biophysical and biogeochemical processes could explain resource transfer in CMNs with more accuracy than previous plant- or fungi-centric analogies. We use a grassland system where fungal ASVs are frequently found within the roots of multiple plants in a small area, a system that meets the broader CNM definition given by Rillig et al (2024). In our study, we focus on dynamics in a system that has a high probability of connectivity. We quantify interspecific carbon and nitrogen transfer focusing on plant traits that are known to regulate physiological performance (Dawson et al., 2022), rather than aiming to prove that CMNs are the only explanation. By measuring plant traits and environmental variables that affect resource-use efficiencies across different species we describe how the transfer of carbon and nitrogen occurs in paired experiments designed to affect soil water and nutrient mass flow. We labeled perennial plants central to each plot (hereafter, ‘donors’) with stable isotopically enriched gases and monitored leaf ^15^N and ^13^C for the surrounding plants (‘receivers’), monitored from immediately after labeling to 21 days after. We replicated our paired experimental setting at three different locations, with study sites distributed across a 520 km latitudinal gradient. We also sequenced strain-level variation in root fungal DNA, plant functional types, and leaf stoichiometric traits to test if relatedness (same or different species as the donor) explained difference in resource transfer. Both plants and fungi have economic spectra characterized by contrasting traits and nutrient strategies which together form an interacting continuum potentially driven by resource use and availability (Ward et al., 2022). It is unclear to what extent plant or fungi characteristics drive these plant-fungal interactions. Therefore, we focused on quantifying how plant-fungal interactions influence the structure and functioning of CMNs across environmental gradients and resource constraints.

## Materials and Methods

We conducted our experiment at three sites situated on a 520 km latitudinal transect that spans three Mediterranean climates: cool, moist (northern site; Tenino, WA) to warm, moist (central site; Eugene, OR) to warm, dry (southern site; Selma, OR). Each circular plot was 3 m in diameter. Half of our plots were restored prairie systems (n = 10 per site) while the other half of the plots had introduced pasture grasses established prior to restoration (n = 10 per site). Restored prairie plots were mowed, raked, received herbicide, and seeded in 2014-2015, followed by seeding in fall 2015, 2016, and 2017 (Reed et al., 2019). We erected rainout shelters that excluded 40% of the rainfall on half the plots at each site (n = 10 rain exclusion, 10 control per site; Fig. 1). Due to the climatically driven differences in communities across sites, not all species were present at all sites (Table S1); however, all functional groups were present at all sites and most species were present at more than one site. Our experimental design was nested in a multi-year experiment where data loggers were used to continuously measure temperature and moisture in all the manipulated plots.

**Figure 1.**
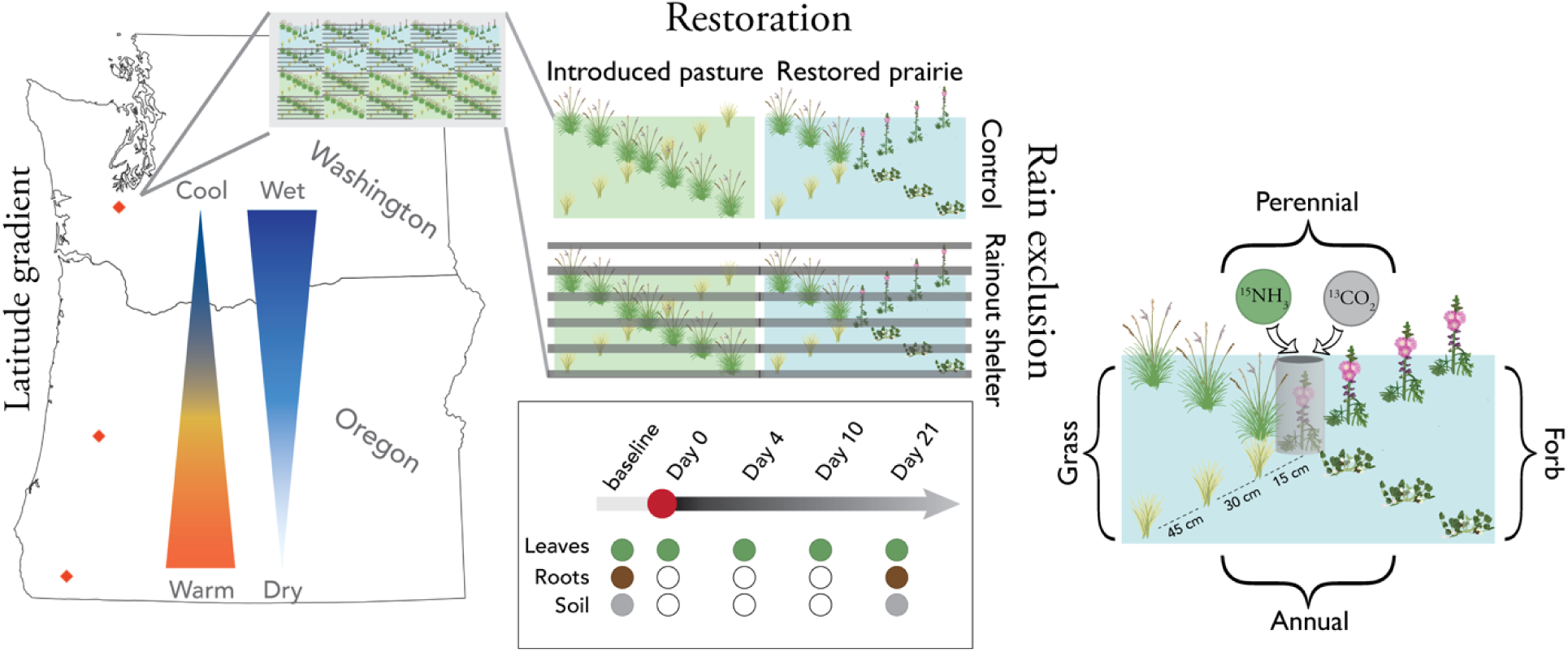
Schema of experimental set up, sampling, and effects of rainout shelters. Figure shows the three location of the sites (orange points on map) in relation to the climatic gradient (cool to warm, wet to dry). Each plot had a central perennial plant that was labeled with isotopically enriched gas (shown with a grey cylinder here) and we sampled plants of each functional type at three distances from the central labelled plant. Inset shows the temporal sampling scheme of leaves, roots and soil.

We recognize that there are many challenges for establishing field experiments of CMN effects, such as treatments for severed versus intact connections (Karst et al., 2023). The plants we experimented on grow in a shared plot where nutrients could transfer via soil, water, bacteria, other fungal guilds, or other non-CMN mechanisms. This does not exclude the possibility that fungi play a large role in these interactions, especially in a system where all plant species have the potential to engage in the most common form of a mycorrhizal connection (Table S1; Heijden et al., 2015). As Rillig et al. (2024) point out, maintaining a strict definition for CMN limits research from exploring meaningful ecological interactions. We have approached this experiment under their proposed broader definition of CMN, an approach which allows us to advance our understanding of how CMNs may function in natural ecosystems.

Previous work demonstrated that the rainout shelters had minimal effects on aboveground community structure or function (Dawson et al., 2022), possibly due to the shoulder season effect of the Mediterranean rain seasonality. This network of experimental sites was established in 2010 and has been extensively studied since then (Brambila et al., 2023; DeMarche et al., 2021; Reed, Bridgham, et al., 2021; Reed et al., 2019, 2023; Reed, Peterson, et al., 2021; Reed, Pfeifer■Meister, et al., 2021), including work on mycorrhizal fungi (Vandegrift et al., 2015; Wilson et al., 2016). Treatments had marginal effects on the soil water potential (especially during the early growing season). Despite those differences, there were no significant changes in the plant community composition or productivity under rain exclusion, which also did not affect morphological and functional traits (i.e., specific leaf area, iWUE, and C:N ratios) of the functional groups we selected for this experiment (Dawson et al., 2022; Reed, Pfeifer■Meister, et al., 2021).

### Isotopic labelling

At each site, we selected a healthy perennial forb (*Sidalcea malviflora* ssp*. virgata* in restored prairie plots [except in one plot where we used *Eriophyllum lanatum* due to a lack of *S. malviflora ssp. virgata*]), or a perennial grass (*Alopecurus pratensis, Schedonorus arundinaceus,* or *Agrostis capillaris*) in pasture plots at the center of each plot to receive the isotopic labels. On sunny days between 11AM and 3PM, we applied isotopically enriched carbon (^13^C) and nitrogen (^15^N) as a pulse of carbon dioxide (CO_2_) and ammonia (NH_3_) to the leaves of target “donor” species common across experimental sites. Although gases are not the primary source of nitrogen for most plants, applying gaseous nitrogen allowed us to limit the amount leaked into the soil compared to applying nitrogen directly to the soil (Silva et al., 2015). Plant leaves are known to uptake ammonia (Farquhar et al., 1980; Sutton et al., 2008). We performed the labeling experiment using custom-made clear chambers with internal fans, following established protocols (e.g., Earles et al., 2016; Silva et al., 2015; Sperling et al., 2017). Before performing the experiment, we tested our approach in the field to optimize gas exposure and labeling amounts, which included checking for leaks and contamination outside of the chamber. We covered the donor plant with a clear plastic cylindrical chamber and injected gas in sequence at 20 minute intervals. For ^13^CO_2_, we made three injections of 2 mL pure CO_2_ (98 atm % ^13^C) to double the amount of CO_2_ in the chamber each time. For NH_3_, we made two injections of 10 mL pure NH_3_ (98 atm% ^15^N). The dates of application were based on peak productivity estimated from Normalized Different Vegetation Index (NDVI) at each site (see Reed et al., 2019 for details). We sampled leaves from each donor plant immediately after labeling (time point 0) as well as from all plants approximately 4 days (time point 1), 10 days (time point 2), and 21 days (time point 3) post-labelling (Fig. 1, Table S2). Time points were chosen to balance the potentially rapid transfer of nutrients through the system with the logistical difficulties of a single team sampling along a 520 km gradient. We also collected leaves at time points 1, 2, and 3 from up to twelve plants in each plot representing three replicates of grass/forb structural groups and annual/perennial life history strategies (Table S1). The number of plants and groups depended on which plants were growing in each plot.

At the end of the experiment, we harvested entire plants and the soil surrounding the roots at time point 3 and kept them in cool conditions until processing. We separated the roots and rhizosphere soils, and selected approximately ten ∼3 cm fine root fragments per sample (i.e., third order or finer, where available) for DNA extraction and identification. All roots and rhizosphere soils were stored at −80° C until processing.

### Baseline and Resource Transfer Calculations

Before isotopic labeling, we collected soil, leaves, and roots from each site. We collected soils in late spring and early summer 2019 to 20 cm depth in each plot. From these soil samples, we removed root fragments that represented the typical roots seen in each plot. We collected leaves for each species in each plot; however, these leaves were contaminated with ^15^N during transport from the highly enriched time point 0 donor samples that were transported with them. To replace contaminated samples, we separately sampled leaves from biomass samplings collected in late spring and early summer 2019, ensuring that annual and perennial grasses and forbs were represented at each site.

We oven-dried all samples at 65° C to constant mass and encapsulated them for stable isotope analysis. All stable isotope analysis was done at UC Davis Stable Isotope Facilities. We calculated the amount of carbon and nitrogen in each plant compartment (leaves and roots) using standard label recovery equations (Silva et al., 2015), using baseline values measured before application of the labelled gases to capture background variations in isotopic composition of unenriched leaves, roots, and soil samples.

We designated all samples with greater than two standard deviations above baseline samples as “enriched” in a particular isotope. Baseline values were calculated on a site by rain exclusion treatment basis by plant functional type basis (Table S3). Site-specific baseline soil and root isotope ratios represent the whole community because we were unable to differentiate which roots belonged to which plants from our soil cores.. In all cases, baseline values fell within the expected range for our region (Fig. S1). For each enriched sample, we calculated isotope excess as

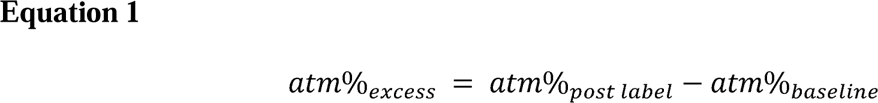

For enriched donor plants, we calculated % derived from label immediately following label application as

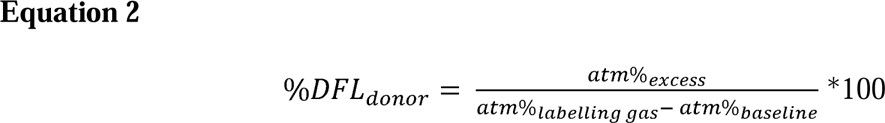

For enriched receiver plants, we calculated % derived from label (%NDFL and %CDFL, respectively) for each relevant point in space and time as

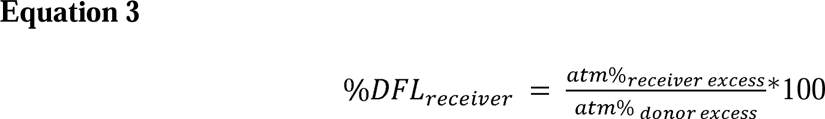

When we calculated %DFL in roots, we used donor leaves as the source (atm% donor excess). We then calculated the amount derived from label on a per mass basis as

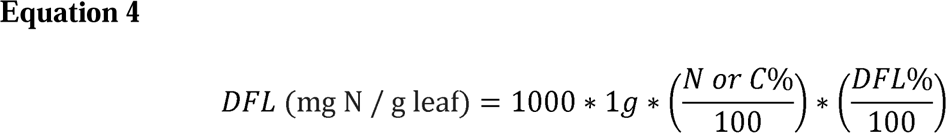

We calculated intrinsic water-use efficiency following Farquhar and Richards (1984) as detailed in (Dawson et al., 2022) using the baseline ^13^C values from the original baseline samples. The equations given in Farquhar and Richards (1984) scale δ13C of plant tissue samples with atmospheric data and by plant physiological constants. Because of sampling discrepancies, we did not have intrinsic water-use efficiency data for 32 plants.

We selected a subset of rhizosphere soils that represented six donor plants at each site divided equally between restored prairie and pasture plots, and selected the three most highly ^15^N-enriched interspecific receivers in each plot across the sites. In addition, we sampled three most highly enriched interspecific receivers at each site and restored prairie-introduced pasture combination. We sampled the top three enriched intraspecific receivers at each site and treatment. In total, this came to 48 post-labelling soil samples in 29 plots.

### Fungal DNA analysis

We extracted DNA from roots of 450 plants harvested at time point 3 (21 days post-label) using Qiagen DNeasy Powersoil HTP kits (Qiagen, Hilden, Germany). We only analyzed DNA from roots, not from the soils collected from each plant’s rhizosphere. We characterized each sample’s AM fungal composition with a two-step PCR protocol that amplified a ∼550bp fragment of the SSU rRNA gene (the most well-supported region for AM fungal taxonomic resolution (Dumbrell et al., 2011). We used WANDA (5’-CAGCCGCGGTAATTCCAGCT-3’) and AML2 (5’-GAACCCAAACACTTTGGTTTCC-3’) primers (Langmead & Salzberg, 2012; Lee et al., 2008). We used primers with unique indices so we could multiplex several projects on a single run. We quantified successful PCR amplicons with the Quant-iT PicoGreen dsDNA Assay Kit (Invitrogen, Waltham, MA, USA) on a SpectraMax M5E Microplate Reader (Molecular Devices, San Jose, CA, USA) before purifying with QIAquick PCR Purification kits (Qiagen). We sequenced the purified pools on the Illumina MiSeq platform (paired-end 300bp, Illumina Inc., San Diego, CA, USA) at the University of Oregon Genomics and Cell Characterization Core Facility (Eugene, OR, USA). Reads were deduplicated with UMI-tools using unique molecular identifiers (UMIs) inserted during PCR (Smith et al., 2017).

We assigned amplicon sequence variants (ASVs) using the dada2 pipeline (version 1.18.0) with standard quality filtering and denoising parameters (Callahan et al., 2016). The dada2 pipeline maintains strain-level diversity at the scale of individual sequence variants rather than clustering sequences into OTUs. This fine-scale measure of fungal sequence diversity was particularly important for our analyses to maintain the greatest chance of detecting a single AM fungal ‘individual’ in multiple plant root samples. Taxonomy was assigned to ASVs using the MaarjAM database (2019 release) (Öpik et al., 2010). We used a Bayesian mixture model in the DESeq2 package (Love et al., 2014) to scale ASV counts within and across samples to avoid artificial taxon abundance biases (Anders & Huber, 2010).

### Replication statement

| Scale of inference | Scale at which the factor of interest is applied | Number of replicates at the appropriate scale |
| --- | --- | --- |
| Individuals | Individual donors and receivers | 1,441 leaves, 450 root DNA samples, and 364 root stable isotope samples from 60 donor and 502 receiver plants |
| Species | Functional groups | 18 species with at least 3 individual plant replicates each (details in Table S1) Grouped by 61 annual forb plants, 131 annual grass plants, 81 perennial forb plants, 161 perennial grass plants |
| Community | Pooled by site and<br>by climate and restoration treatment | 3 sites (northern, central, and southern latitude sites spanning ~520 km; 20 plots per site; 60 total)<br>Rain exclusion (10 plots per site; 30 total) vs ambient (10 control plot per site; 30 total); Restored prairie (10 plots per site; 30 total) vs pasture (10 plots per site; 30 total) treatments; |

### Data analysis

We performed all analyses in R ver. 4.0.4 (R Core Team, 2022). Graphs were made in ggplot2 (Wickham, 2016). We removed one outlier plant with a ^15^N atm% more than twice as high as the next highest measurement. We also removed five mislabeled samples. To meet statistical assumptions, we only included data from enriched plants with successful root fungal DNA extraction in our analyses and figures. We limited receivers to those for which we also had sufficient root fungal DNA. In total, from 1441 leaves measured for isotopic content and with successfully recovered fungal DNA, we analyzed data from 353 unique plants: 54 donors and 353 receivers. We excluded two plots (one central rain exclusion restored plot and one central rain exclusion pasture plot) because either no donor or no receiver leaves were recovered.

We tested the relationship between receiver leaf %DFL and plant traits (grass/forb, annual/perennial, iWUE, C:N, degrees of connectivity, interaction term between grass/forb and annual/perennial) and site conditions (position on latitude gradient, pasture/restored, rain exclusion treatment, distance from donor, time from labelling) with a mixed-effect ANOVA (plot nested within site as random effect; Table 1). We used a Tukey post-hoc test for differences within groups shown in the following figures and tables. We constructed a phyloseq object using the ASV table with normalized counts (McMurdie & Holmes, 2013), and used iGraph, metagMisc, and RCy3 (Gustavsen et al., 2019; Mikryukov, 2017; Nepusz & Csardi, 2006) to create networks for each plot. In each network, nodes represented individual plants and edges between nodes represent plants sharing at least one fungal DNA sequence variant. The weighted edges are based on how many fungal ASVs were shared among plants. We calculated degrees of connectivity with tidygraph (Petersen, 2022) to examine how many plants each individual plant was ‘connected’ to (by means of shared fungal ASVs) in each plot (Fig. S2). We also calculated whether each receiver plant shared fungal ASVs with the central donor plant in each plot. We visualized shifts in AM fungal community composition using non-metric multi-dimensional scaling (NMDS) in the vegan package, demonstrating the AM fungal community similarity across plants (Oksanen, et al., 2022).

**Table 1.**
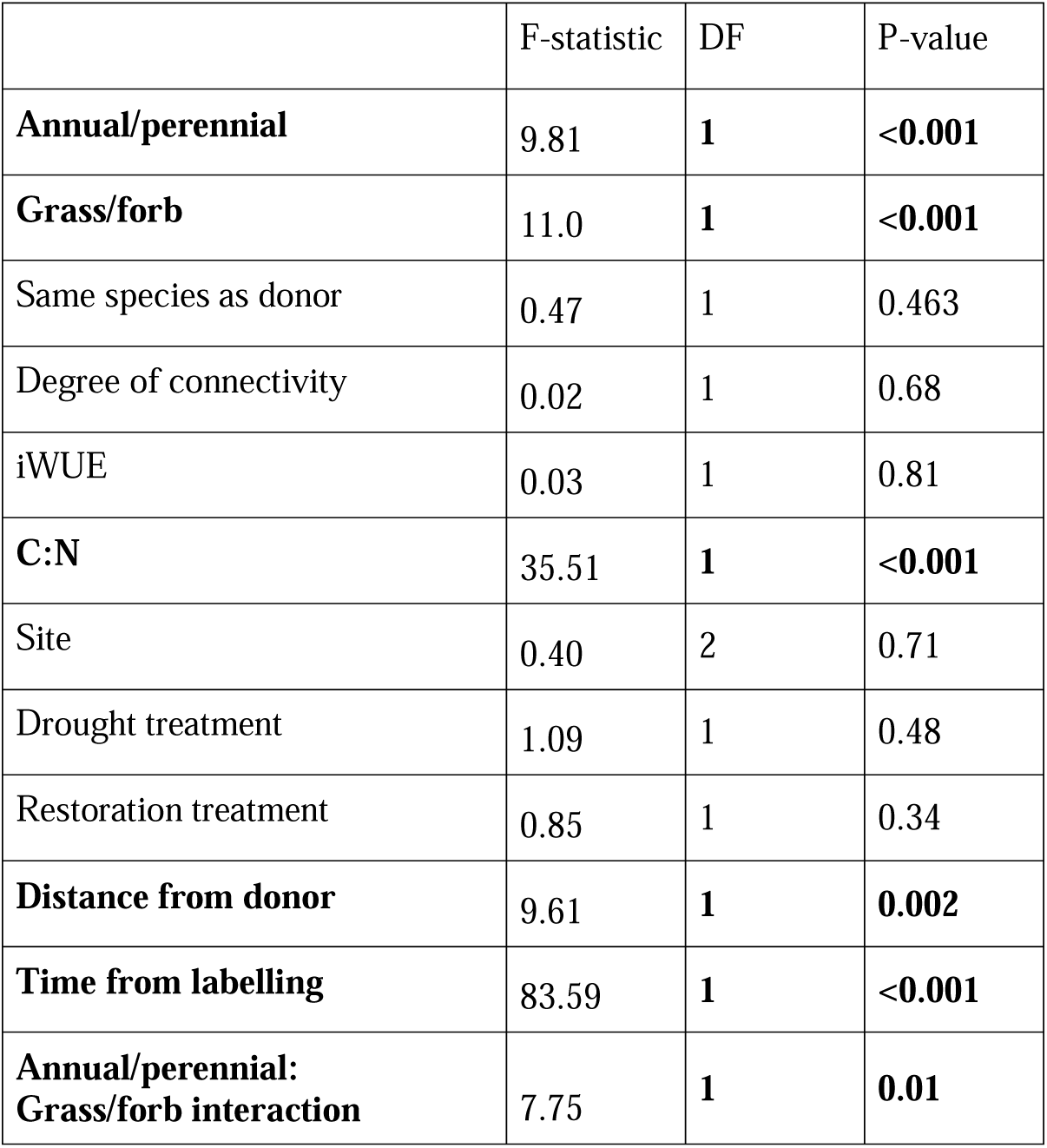
Mixed-effects ANOVA results effects on leaf nitrogen derived from label (%NDFL) Random effect is plot nested within site. Only enriched receiver leaves with associated DNA data were included in the analysis (n = 1094). Leaf %CDFL results available in Table S4.

|  | F-statistic | DF | P-value |
| --- | --- | --- | --- |
| **Annual/perennial** | 9.81 | **1** | **<0.001** |
| **Grass/forb** | 11.0 | **1** | **<0.001** |
| Same species as donor | 0.47 | 1 | 0.463 |
| Degree of connectivity | 0.02 | 1 | 0.68 |
| iWUE | 0.03 | 1 | 0.81 |
| **C:N** | 35.51 | **1** | **<0.001** |
| Site | 0.40 | 2 | 0.71 |
| Drought treatment | 1.09 | 1 | 0.48 |
| Restoration treatment | 0.85 | 1 | 0.34 |
| **Distance from donor** | 9.61 | **1** | **0.002** |
| **Time from labelling** | 83.59 | **1** | **<0.001** |
| **Annual/perennial:**<br>**Grass/forb interaction** | 7.75 | **1** | **0.01** |

## Results

All donors were ^15^N-enriched in their leaves at time of labelling. Two donors were not ^13^C-enriched in their leaves at time of labelling. We recovered DNA data from the roots of 88.3% of donors. We sampled 1,444 leaves from 434 receiver plants at three time points. Of these leaves, 81.0% were ^15^N-enriched and 2.4% were ^13^C-enriched. We recovered DNA from the roots of 77.9% of the receivers. At time point 3 (∼21 days post-labelling), we collected roots from 46 of the initial 60 donors and 306 of the initial 434 receivers. Of the roots of the collected donors, 97.8% were still 15N-enriched and 23.9% were still ^13^C-enriched at time point 3. Of the roots of the collected receivers, 33.0% were still ^15^N-enriched and 10.5% were still ^13^C-enriched at time point 3.

Assimilation of isotopic tracers was similar between labeled “donor” plants with no significant differences on average between sites or experimental treatments within sites, including rainfall exclusion or restored status (Fig. S3). At all sites, foliar assimilation of ^15^N and ^13^C by donor plants led to enrichment levels ranging from approximately 5-10 fold higher than baselines. Foliar enrichment levels decreased consistently at all sites and treatments over the 21 day sampling period. Annual forbs had the greatest enrichment level and perennial forbs the lowest enrichment level (Fig. 2). We found significant spatial and temporal differences in foliar and root isotope ratios in donors and receivers resulting from interspecific transfer of carbon and nitrogen (Fig. S4; Table 1). Receiver foliar enrichment levels did not correlate with donor foliar enrichment levels within the same plot (Fig. S5).

**Figure 2.**
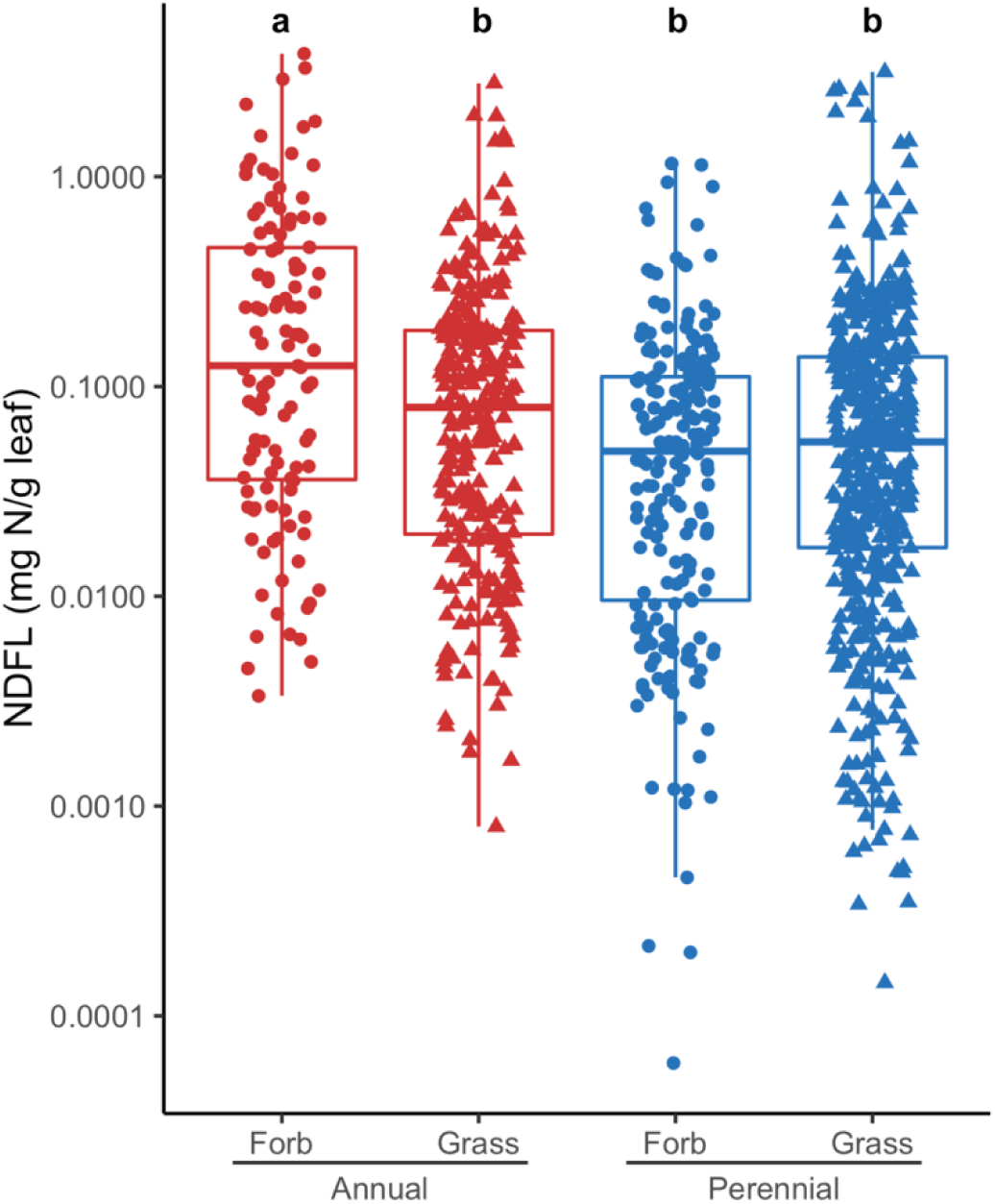
Annuals had greater leaf nitrogen derived from label (NDFL) scaled by mass than perennials. Points represent enriched individual receiver plants with associated DNA data at one time point of sampling. Y-axis is log10 scale with raw value datapoints. Boxplots show medians and interquartile ranges, lines show the largest (or smallest) value no further than 1.5x IQR from the hinge. Letters above the boxplots indicate significant differences.

Allocation and transfer varied significantly between functional groups due to their intrinsic differences in tissue stoichiometry (Table 1). We selected 18 common annual/perennial and grass/forb species of receiver plants, which revealed significant differences between functional types for NDFL (Fig. 2) but little detectable CDFL relative to baseline (Fig. S4, S6). Rain exclusion treatment, restoration treatment, and site did not affect interspecific transfer of nitrogen (Fig. S7; ANOVA, P > 0.05, Table 1). We observed very low carbon enrichment, but of the 2.4% of leaves that were enriched in carbon, plant functional type and site affected carbon transfer (Table S4). We did, however, observe significant differences in nitrogen transfer by plant functional type (Table 1), mirroring intrinsic differences in tissue stoichiometry and iWUE (Fig. 3), despite no significant enrichment in soils collected from the rhizosphere of those same plants (Fig. S8). We also found that C:N affected NDFL, although not in a simple linear manner and with no apparent correlation between NDFL and iWUE (Fig. 3). We did detect a low level of soil enrichment in 1 out of 30 receiver soil samples (0.377 atm% ^15^N) and 5 out of 18 donor soil samples (ranging from 0.376 to 0.479 atm% ^15^N; Fig. S8).

**Figure 3.**
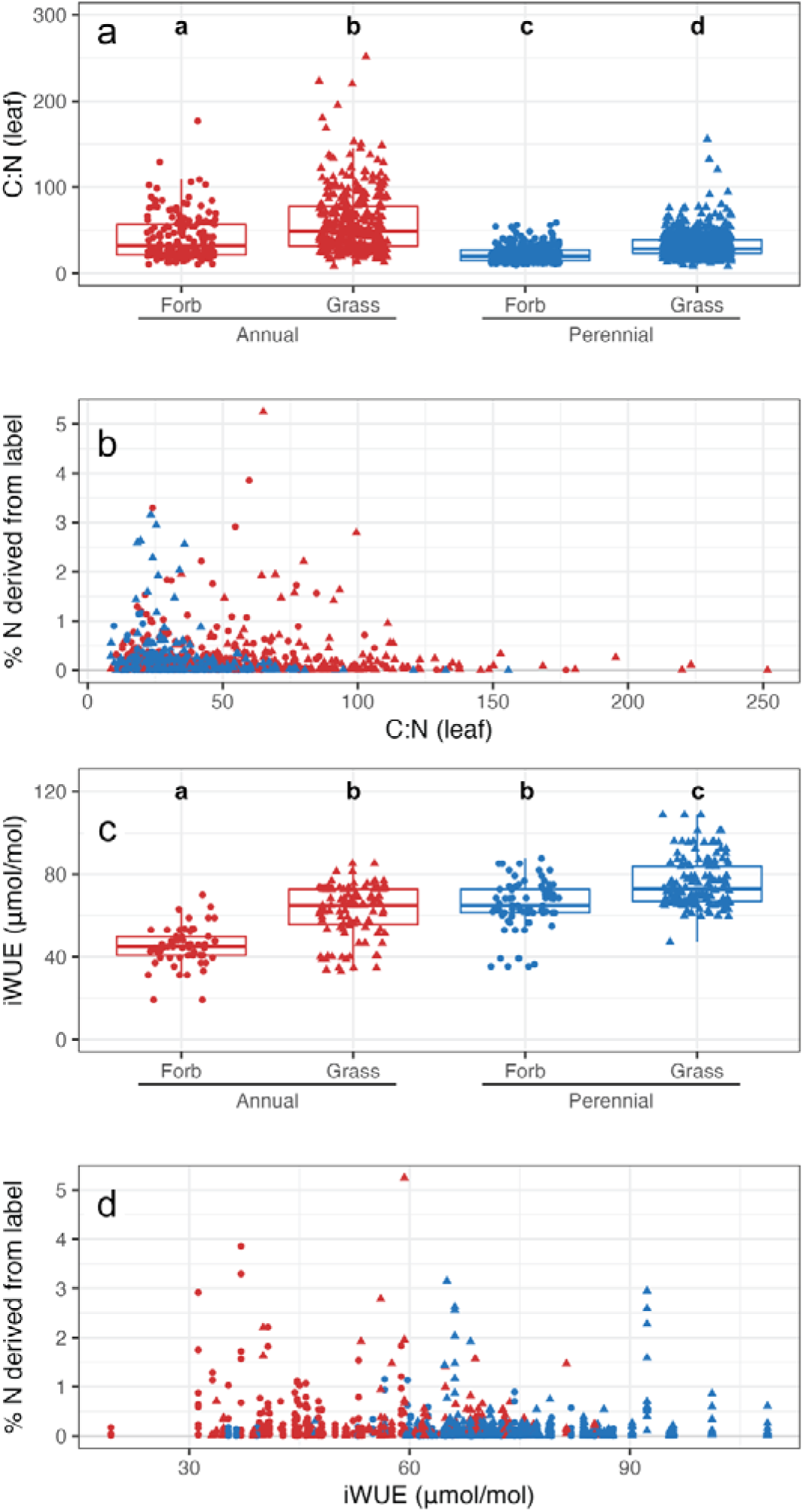
Stoichiometric and functional traits compared to nitrogen derived from transfer. A) Leaf C:N by annual/perennial. B) Percent ^15^N derived from label (DFL) compared to C:N in leaves. C) iWUE by annual/perennial as measured before labelling. D) Percent ^15^NDFL compared to iWUE in leaves at all time points. Boxplots show medians and interquartile ranges, lines show the largest (or smallest) value no further than 1.5x IQR from the hinge. Letters above boxplots indicate significant differences.

Annuals had greater ^15^N foliar enrichment compared to perennials (ANOVA, P < 0.001, Tables 1 and 2). Foliar enrichment decreased over both time and space (ANOVA P < 0.001; Fig. S3, Fig. S4). On average, annuals had a lower leaf nitrogen content and higher C:N than perennials (Table 2, Fig. 3). Forbs had higher NDFL than grasses (ANOVA, P < 0.001, Table 1) as well as a lower C:N. There was a significant interaction between annual/perennial and grass/forb form (ANOVA, P = 0.003, Table 1). There was no correlation in ^15^N-enrichment and whether the donors and receivers were the same species (Table 1).

**Table 2.** Leaf nitrogen derived from label (NDFL) and leaf tissue nitrogen (N%) four days after labeling.

|  |  | *n* | %NDFL |  |  |  | NDFL (mg N/g leaf) |  |  |  | N (%) |  |  |  |
| --- | --- | --- | --- | --- | --- | --- | --- | --- | --- | --- | --- | --- | --- | --- |
|  |  |  | *Mean* | *SD* | *Min* | *Max* | *Mean* | *SD* | *Min* | *Max* | *Mean* | *SD* | *Min* | *Max* |
| **Annual** | **Forb** | 54 | 6.528 | 9.769 | 0.101 | 56.005 | 0.749 | 0.819 | 0.041 | 3.858 | 0.749 | 0.819 | 0.041 | 3.858 |
|  | **Grass** | 92 | 4.847 | 9.032 | 0.098 | 63.884 | 0.34 | 0.463 | 0.011 | 2.79 | 0.340 | 0.463 | 0.011 | 2.790 |
| **Perennial** | **Forb** | 67 | 0.952 | 1.069 | 0.090 | 5.582 | 0.214 | 0.228 | 0.021 | 1.156 | 0.214 | 0.228 | 0.021 | 1.156 |
|  | **Grass** | 148 | 1.632 | 3.066 | 0.088 | 22.434 | 0.252 | 0.484 | 0.013 | 2.948 | 0.252 | 0.484 | 0.013 | 2.948 |

Fungal community composition demonstrated a high degree of potential connectivity between plants of different species but no obvious pattern of connectedness that could explain preferential nutrient transfer by plant functional groups. We found that 97.25% ± 8.01 (SD) of all plant roots within each experimental plot shared at least one fungal DNA sequence variant (ASV) with another plant of the same plot. Fungal community composition was similar across plant functional groups (Fig. 4, PERMANOVA pseudo-F statistic = 2.269, R^2^ = 0.005, p = 0.001). Annual plants shared fungi with more plants in the same plot (4.74 plants ± 2.62 SD) compared to perennials (4.04 plants ± 2.49 SD; t-test, P = 0.019; Table S5), but degrees of connectivity did not predict nitrogen transfer (Table 1). Seventy-three percent of plants were colonized by four or fewer fungi and shared fungi with five or fewer other plants in the plot, making it difficult to determine if strength of connectivity altered nitrogen transfer (Fig. S9).

**Figure 4.**
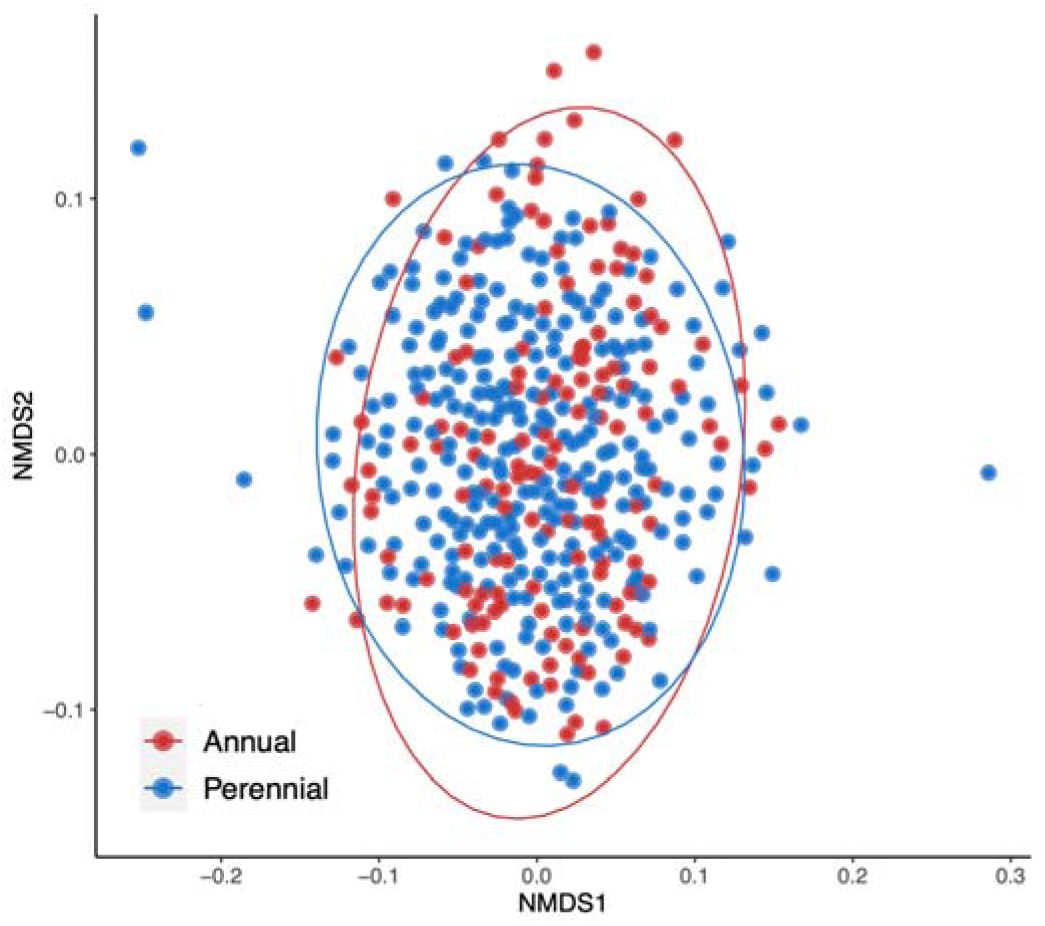
NMDS of fungal communities. Each point represents an individual plant ordinated by the Bray Curtis dissimilarities for AM fungal composition. PERMANOVA pseudo-F statistic = 2.269, R^2^ = 0.005, p = 0.001.

## Discussion

In a system where fungal DNA sequence variants were shared between ∼97% of plants, we found that the transfer of nitrogen is regulated by plant functional traits that are known to influence resource use and allocation in plant communities. Using a recently proposed definition of CMNs that prioritizes ecological understanding (Rillig et al., 2024), our data suggest that the assimilation and allocation of limiting resources in CMNs was neither plant-nor fungi-centric. In our study, the rates and direction of resource transfer in our potential CMNs, inferred from pulse labeling and recovery of ^15^N in leaves and rhizospheres, could be predicted from leaf C:N and distance from donor species (Table 1). Across all sites and treatments, we observed a stronger sink for ^15^N in annual plants (Table 2), indicating preferential transfer of limiting resources to that functional group of plants. We applied the ^15^N enriched label to perennial species in all plots, so this greater enrichment in annual plants precludes a preference for receiver plants of the same species as would be expected under the kinship hypothesis (Table 1). Although nearly all plants shared fungal ASVs in their roots (Fig. 4), connectivity did not predict ^15^N transfer (Table 1). However, we did not test hyphal continuity of our network and we draw our conclusions based on an ecological definition of CMNs, not under Rillig et al.’s (2024) definition of CMN-HC. Our data suggest that rates and direction of resource transfer in CMNs reflect plant nutrient requirements and spatial proximity.

We conducted repeated spatiotemporal sampling of isotopic-enrichment levels at increasing distances from donor species, days to weeks after labelling, and in well-established communities exposed to multiple years of experimental treatments, expecting to find evidence of kinship (i.e., greatest resource transfer in plants of the same species), driven by CMN economics (i.e., ^15^N transfer rates coinciding with ^13^C investment in root and fungal mass). Annuals received, on average, an order of magnitude higher enrichment than perennials even though our donor plants were perennials. We also did not find evidence of a relationship between ^15^N in leaves and ^13^C in roots because we did detect ^15^N-enrichment in leaves but no ^13^C-enrichment in roots. We did, however, find significant ^13^C-enrichment in the donor plants. Therefore, our data do not support either hypothesis, and instead suggest AM fungi form CMNs where the rates and direction of resource transfer ultimately reflects a sink-source strength effect, consistent with previous observations of stoichiometric source-sink manipulations of carbon and nitrogen within plants (Cai et al., 2021; Ruiz-Vera et al., 2017; Tegeder & Masclaux■Daubresse, 2018), but in our case observed at the community scale.

Nitrogen enrichment levels remained high in leaves and many roots at the end of the experiment, allowing us to measure NDFL across the community and infer the main predictors of N transfer. However, carbon enrichment levels faded before plants were harvested approximately 21 days post-labelling. After controlling for variation in assimilation rates by calculating NDFL, we found that annual plants received greater ^15^N-enrichment than perennial plants. Plants closest to the donor were most enriched, and ^15^N-enrichment decreased over time (Fig. S4, Fig. S5). Although the rainout shelters had limited effect, there were major differences across the latitudinal gradient represented by the sites in temperature and soil moisture availability (Dawson et al., 2022; Reed et al., 2019); however, neither treatment nor site affected our results.

The major predictors of differences in allocation of ^13^C and ^15^N to roots and subsequent transfer to “receiver” species were the intrinsic difference in plant functional types and correlated traits, including measured leaf C:N. Previous studies in northern California under environmental conditions similar to those found in our southernmost experimental site showed rapid (days to weeks) transfer of ^15^N applied to the leaves of ectomycorrhizal pines to surrounding annual AM plant receivers (He et al., 2006). Those results demonstrated “direct fungal connections are not necessary for N transfer among plants” and, similar to our results, the “leaves of the annual plants had greater ^15^N derived from source and were more enriched (^15^N at % excess and δ^15^N values) than perennial receivers, irrespective of the mycorrhizal type.” Similarly, as proposed by He et al. (2006), our data suggest that annual plants were a strong sink for N which could be explained by stoichiometric gradients that affect root exudation and recapture of N-containing materials from rhizodeposition (Høgh-Jensen & Schjoerring, 2001; Mayer et al., 2003). Although our study included two species with symbiotic N fixation ability that could have altered some of the baseline data, even if a plot was fully dominated by legumes that difference would represent a minor effect relative to the pulse label application. Our data corroborate rapid transfer among AM plants, with no detectable enrichment in root or soil ^13^C near roots 21 days post-labelling, but do not allow us to determine general mechanisms that are responsible for the ^15^N transfers. Other methods of transfer–such as by fungi of other functional guilds, bacteria, or water flow in the soil–were possible given that plants shared a common growing medium in each plot.

We inferred a high connectivity between plants within each given treatment and site given the highly similar fungal composition in the root systems of both perennial and annual plants (Fig. 4). Given the constraints of ASV-identified data, we did this analysis on a strain-level scale and it is possible that separate spores of the same ASV separately infected plants within the same plot. However, we are reasonably confident in our use of fungal ASVs as a proxy for connectivity given the strong overlap in our community and because individuals of one ASV can anastomose in the soil (Mikkelsen et al., 2008). This overlap could explain the lack of support for the kinship hypothesis in our dataset and offers further support for stoichiometric gradients in general, and C:N gradients in particular, as a principal control of terms of trade in CMNs (Kiers et al., 2011). Because we observed such high rates of shared ASVs (∼97%) and we observed high levels of N-enrichment in receivers at our first post-label sampling point four days after application, we could not test whether receivers connected to donors or whether plants were connected to the network affected ^15^N transfer. We found unexpectedly low soil isotopic-enrichment (fig. S8) which suggested that the labels did not remain in the fungal network. This low level was likely driven by the fact that we did not sample soils until 21 days after labeling and hyphal turnover for grass-associated AM fungi can be less than one month (See et al., 2022). Because N is a major limiting nutrient in this system, according to the current paradigm of CMNs, it would be quickly taken up and recycled or transferred rather than accumulating in the soil.

We found that plant-soil stoichiometric gradients and functional traits were the strongest predictors of resource sharing in a possible grassland arbuscular mycorrhizal CMN. We interpret this finding as evidence of biochemical and biophysical sinks, in which nutrients are allocated to plants with the greatest need for those nutrients, either through a ‘passive’ mycorrhizal network as suggested by the high number of shared fungal ASVs or direct uptake from soil or water flow. Expanding on previous studies, we propose that AM fungi facilitate spatiotemporal dynamics of carbon and nitrogen transfer through CMNs in ways that are neither plant-nor fungi-centric. That is, plants and fungi that are located closer together in space and with stronger demand for resources over time are more likely to receive larger amounts of those limiting resources.

