## Supplemental Information for "Plant functional types and tissue stoichiometry explain nutrient transfer in common arbuscular mycorrhizal networks of temperate grasslands"

**Table S1. List of species at each site with number of leaf samples per species.** Donor species are indicated in bold.

| | | | Site | | | | Mean ± SD stoichiometry | |
|---|---|---|---|---|---|---|---|---|
| **Species** | **Annual/ perennial** | **Grass/ forb** | **Northern** | **Central** | **Southern** | **Fungal symbiont** | **N** | **C** |
| ***Agrostis capillaris* L.** | Perennial | Grass | 121 | -- | -- | AM[1] | 1.34 ± 0.71 | 41.02 ± 4.79 |
| ***Alopecurus pratensis* L.** | Perennial | Grass | -- | 210 | -- | AM[1] | 1.3 ± 0.46 | 39.77 ± 6.48 |
| *Aphanes occidentalis* L. | Annual | Forb | 3 | -- | -- | AM*[2] | 0.95 ± 0.47 | 41.41 ± 3.72 |
| *Bromus diandrus* Roth | Annual | Grass | 49 | -- | -- | AM*[1] | 1.06 ± 0.56 | 40.05 ± 7.25 |
| *Bromus hordeaceus* L. | Annual | Grass | 35 | 42 | 96 | AM*[1] | 1.04 ± 0.67 | 37.66 ± 7.62 |
| *Elymus repens* (L.) Gould | Perennial | Grass | -- | -- | 63 | AM*[1] | 1.67 ± 0.82 | 35.8 ± 8.75 |
| *Eriophyllum lanatum* (Pursh) Forbes | Perennial | Forb | 8 | -- | -- | AM[2] | 1.15 ± 0.35 | 37.59 ± 8.86 |
| *Festuca idahoensis* Elmer ssp. *roemeri* (Pavlick) S. Aiken | Perennial | Grass | 18 | -- | -- | AM*[1] | 1.08 ± 0.36 | 41.06 ± 2.82 |
| *Geranium dissectum* L. | Annual | Forb | 15 | 81 | -- | AM[2] | 1.03 ± 0.57 | 39.35 ± 5.41 |
| *Holcus lanatus* L. | Perennial | Grass | -- | 6 | -- | AM[1] | 1.13 ± 0.46 | 40.54 ± 4.07 |
| *Koeleria macrantha* (Ledeb.) Schult. | Perennial | Grass | 18 | -- | -- | AM[2] | 1.2 ± 0.46 | 40.16 ± 3.62 |
| *Lotus corniculatus* L. | Perennial | Forb | -- | 9 | -- | AM[1] | 2.98 ± 1.06 | 42.36 ± 2.78 |
| ***Schedonorus arundinaceus* (Schreb.) Dumort.** | Perennial | Grass | -- | -- | 130 | AM[1] | 1.11 ± 0.48 | 35.68 ± 8.12 |
| ***Sidalcea malviflora* ssp. *virgata* (DC.) A. Gray ex Benth.** | Perennial | Forb | 66 | 129 | 119 | AM[3] | 1.75 ± 0.8 | 37.93 ± 5.96 |
| *Trifolium subterraneum* L. | Annual | Forb | 48 | -- | -- | AM[1] | 2 ± 0.78 | 41.32 ± 4.03 |
| *Veronica arvensis* L. | Annual | Forb | 3 | -- | -- | AM[1] | 1.14 ± 0.41 | 39.2 ± 2.8 |
| *Vicia sativa* L. | Annual | Forb | -- | 14 | -- | AM*[1] | 2.23 ± 1.03 | 41.41 ± 1.91 |
| *Vulpia* spp. | Annual | Grass | 54 | 8 | 21 | AM[1] | 0.91 ± 0.44 | 38.81 ± 7.18 |
| | | **Total samples** | 438 | 499 | 429 | | | |

Notes: Fungal symbiont data derived from [1]Chaudhary et al. (2016), [2]Soudzilovskaia et al. (2020), and [3]Dickie et al. (2013). * indicates "to genus," that strategy was extrapolated from a sister species given a lack of data on this species. Nutrient-use strategies include arbuscular mycorrhizal (AM) and nitrogen fixation (N-fixer).

**Table S2. Collection dates for each time point at each site.** Days since labeling given in parentheses after date.

| **Site** | **Label application (time point 0)** | **Time point 1** | **Time point 2** | **Time point 3** |
|---|---|---|---|---|
| ***Ideal days since labeling*** | *0* | *4* | *10* | *21* |
| **Northern** | June 10 | June 14 (4) | June 20 (10) | July 2 (22) |
| **Middle** | May 29 | June 3 (5) | June 11 (13) | June 18 (20) |
| **Southern** | May 9 | May 13 (4) | May 22 (9) | May 30 (21) |

**Table S3. Baseline values used to calculate amount derived from label (DFL).**

| | | **Roots** | | | | | | **Leaves** | | | |
|---|---|---|---|---|---|---|---|---|---|---|---|
| *Site* | *Drought treatment* | *N atm%* | *±SD* | *C atm%* | *±SD* | *Annual/perennial* | *Grass/forb* | *N atm%* | *±SD* | *C atm%* | *±SD* |
| North | Control | 0.370 | 0.001 | 1.073 | 0.001 | Annual | Forb | 0.367 | 0.001 | 1.07 | 0.001 |
| | | | | | | | Grass | 0.367 | 0.000 | 1.071 | 0.003 |
| | | | | | | Perennial | Forb | 0.367 | 0.000 | 1.072 | 0.001 |
| | | | | | | | Grass | 0.368 | 0.000 | 1.075 | 0.001 |
| | Rain exclusion treatment | 0.370 | 0.001 | 1.073 | 0.001 | Annual | Forb | 0.367 | 0.001 | 1.072 | 0.001 |
| | | | | | | | Grass | 0.367 | 0.000 | 1.073 | 0.000 |
| | | | | | | Perennial | Forb | 0.367 | 0.000 | 1.072 | 0.002 |
| | | | | | | | Grass | 0.368 | 0.000 | 1.075 | 0.001 |
| Central | Control | 0.368 | 0.001 | 1.073 | 0.001 | Annual | Forb | 0.367 | 0.001 | 1.071 | 0.001 |
| | | | | | | | Grass | 0.366 | 0.001 | 1.073 | 0.001 |
| | | | | | | Perennial | Forb | 0.368 | 0.003 | 1.071 | 0.001 |
| | | | | | | | Grass | 0.367 | 0.003 | 1.074 | 0.002 |
| | Rain exclusion treatment | 0.368 | 0.001 | 1.073 | 0.001 | Annual | Forb | 0.367 | 0.001 | 1.071 | 0.001 |
| | | | | | | | Grass | 0.367 | 0.001 | 1.072 | 0.001 |
| | | | | | | Perennial | Forb | 0.366 | 0.000 | 1.071 | 0.001 |
| | | | | | | | Grass | 0.366 | 0.000 | 1.075 | 0.002 |
| South | Control | 0.368 | 0.000 | 1.074 | 0.001 | Annual | Forb | 0.366 | 0.000 | 1.071 | 0.000 |
| | | | | | | | Grass | 0.368 | 0.001 | 1.074 | 0.001 |
| | | | | | | Perennial | Forb | 0.367 | 0.000 | 1.074 | 0.001 |
| | | | | | | | Grass | 0.367 | 0.000 | 1.074 | 0.000 |
| | Rain exclusion treatment | 0.368 | 0.001 | 1.074 | 0.001 | Annual | Forb | 0.366 | 0.000 | 1.071 | 0.000 |
| | | | | | | | Grass | 0.367 | 0.001 | 1.074 | 0.002 |
| | | | | | | Perennial | Forb | 0.367 | 0.001 | 1.074 | 0.001 |
| | | | | | | | Grass | 0.367 | 0.000 | 1.074 | 0.001 |

**Table S4. Mixed-effects ANOVA results effects on leaf carbon derived from label (%CDFL).** Random effect is plot nested within site. Only receiver leaves enriched with $^{13}C$ with associated DNA data were included in the analysis (n = 92). Note that since 13C enrichment was limited, not all treatments were replicated when filtered to only enriched leaves and the results here should be treated with caution. Leaf %NDFL results (which are more robust as most leaves were enriched with 15N) are available in Table 1.

| | F-statistic | DF | P-value |
|---|---|---|---|
| **Annual/perennial** | 22.74 | 1 | **<0.001** |
| **Grass/forb** | 12.43 | 1 | **0.001** |
| **Same species as donor** | 11.64 | 1 | **0.001** |
| Degree of connectivity | 1.74 | 1 | 0.192 |
| iWUE | 0.17 | 1 | 0.678 |
| C:N | 2.28 | 1 | 0.136 |
| Site | 3.16 | 2 | 0.064 |
| Drought treatment | 1.27 | 1 | 0.274 |
| Restoration treatment | 2.79 | 1 | 0.107 |
| Distance from donor | 0.00 | 1 | 0.958 |
| Time from labelling | 1.37 | 1 | 0.246 |
| **Annual/perennial: Grass/forb interaction** | NA | 0 | NA |

**Table S5. List of fungal ASVs by annuals and perennials.**

| Fungal taxon | No. of perennial plants assoc. with | No. of annual plants assoc. with |
|---|---|---|
| *Acaulospora sp877* | 13 | 0 |
| *Claroideoglomus sp744* | 20 | 7 |
| *Claroideoglomus sp745* | 18 | 5 |
| *Claroideoglomus sp746* | 18 | 7 |
| *Claroideoglomus sp749* | 14 | 4 |
| *Claroideoglomus sp751* | 25 | 10 |
| *Claroideoglomus sp757* | 11 | 1 |
| *Claroideoglomus sp758* | 11 | 1 |
| *Claroideoglomus sp781* | 18 | 13 |
| *Claroideoglomus sp782* | 59 | 36 |
| *Claroideoglomus sp783* | 38 | 20 |
| *Claroideoglomus sp784* | 47 | 21 |
| *Claroideoglomus sp811* | 19 | 9 |
| *Claroideoglomus sp812* | 12 | 4 |
| *Claroideoglomus sp813* | 15 | 10 |
| *Claroideoglomus sp815* | 21 | 11 |
| *Claroideoglomus sp816* | 19 | 7 |
| *Claroideoglomus sp817* | 30 | 8 |
| *Glomus sp1024* | 11 | 1 |
| *Glomus sp1025* | 12 | 4 |
| *Glomus sp1029* | 23 | 4 |
| *Glomus sp1031* | 20 | 7 |
| *Glomus sp104* | 12 | 3 |
| *Glomus sp113* | 23 | 10 |
| *Glomus sp114* | 6 | 16 |
| *Glomus sp115* | 21 | 15 |
| *Glomus sp1349* | 13 | 8 |
| *Glomus sp1351* | 18 | 8 |
| *Glomus sp1352* | 15 | 5 |
| *Glomus sp148* | 22 | 11 |

| Fungal taxon | No. of perennial plants assoc. with | No. of annual plants assoc. with |
|---|---|---|
| *Glomus sp149* | 12 | 6 |
| *Glomus sp150* | 27 | 14 |
| *Glomus sp168* | 21 | 12 |
| *Glomus sp169* | 13 | 6 |
| *Glomus sp170* | 15 | 6 |
| *Glomus sp177* | 11 | 1 |
| *Glomus sp191* | 14 | 10 |
| *Glomus sp470* | 15 | 6 |
| *Glomus sp472* | 14 | 4 |
| *Glomus sp488* | 13 | 2 |
| *Glomus sp539* | 13 | 3 |
| *Glomus sp541* | 19 | 3 |
| *Glomus sp565* | 15 | 7 |
| *Glomus sp566* | 12 | 1 |
| *Glomus sp612* | 13 | 3 |
| *Glomus sp614* | 11 | 6 |
| *Glomus sp648* | 12 | 2 |
| *Glomus sp651* | 25 | 10 |
| *Glomus sp661* | 11 | 4 |
| *Glomus sp662* | 14 | 3 |
| *Glomus sp663* | 12 | 8 |
| *Glomus sp667* | 13 | 1 |
| *Glomus sp668* | 14 | 3 |
| *Glomus sp669* | 14 | 5 |
| *Glomus sp670* | 20 | 2 |
| *Glomus sp692* | 14 | 1 |
| *Glomus sp700* | 11 | 2 |
| *Glomus sp701* | 12 | 12 |
| *Glomus sp703* | 19 | 13 |
| *Glomus sp83* | 11 | 0 |
| *Glomus sp85* | 12 | 3 |

| **Fungal taxon** | **No. of perennial plants assoc. with** | **No. of annual plants assoc. with** |
|---|---|---|
| *Glomus sp904* | 18 | 3 |
| *Glomus sp905* | 11 | 2 |
| *Glomus sp906* | 17 | 6 |
| *Glomus sp917* | 29 | 12 |
| *Glomus sp918* | 85 | 48 |
| *Glomus sp919* | 45 | 32 |
| *Glomus sp92* | 11 | 0 |
| *Glomus sp920* | 53 | 33 |
| *Glomus sp921* | 17 | 3 |
| *Glomus sp938* | 14 | 8 |
| *Glomus sp986* | 12 | 1 |
| *Paraglomus sp301* | 19 | 4 |
| *Paraglomus sp302* | 26 | 5 |
| *Paraglomus sp303* | 15 | 4 |
| *Paraglomus sp304* | 25 | 5 |
| *Scutellospora sp1080* | 12 | 6 |
| *Unknown sp21* | 12 | 4 |

Table only includes fungal species associated with at least 10 plants.

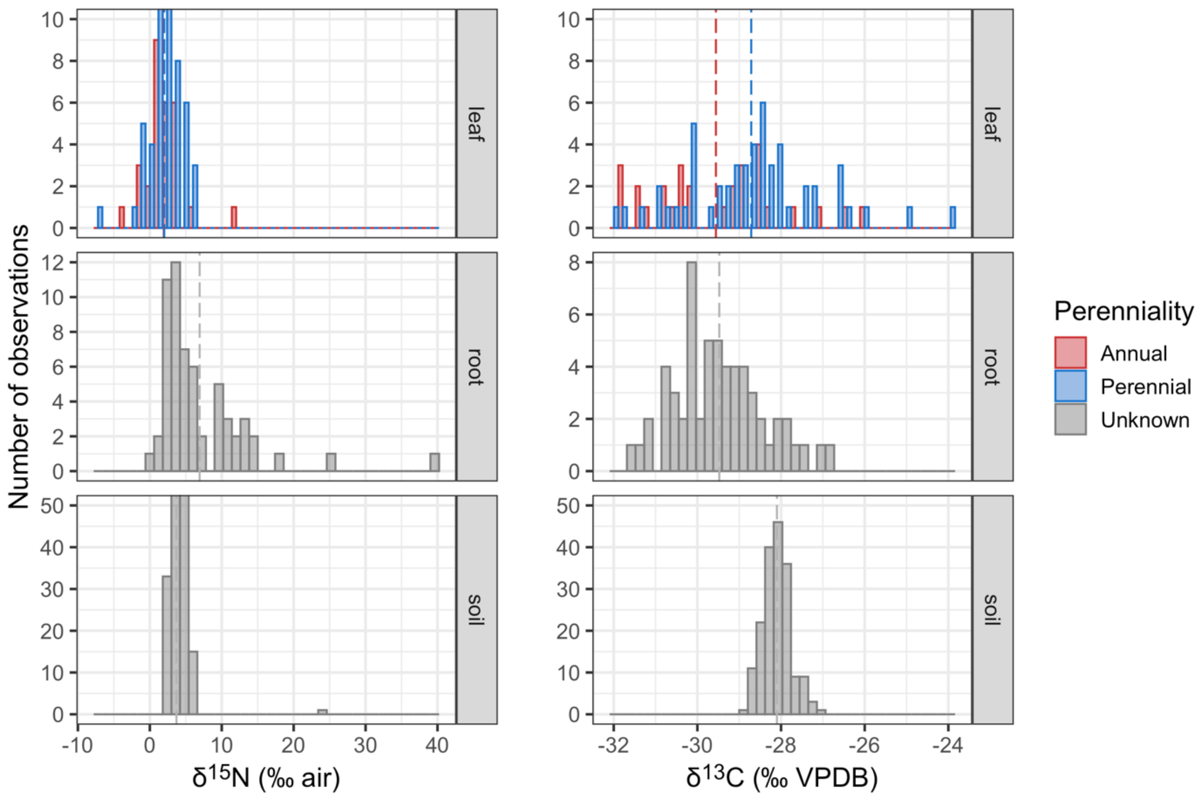


**Figure S1. Natural abundance of stable isotopes in leaves, roots, and soil before labelling.** Dashed lines indicate mean value. Exact values used to calculate derivation from label (Eq. 1 – Eq. 4) are available in Table S3.

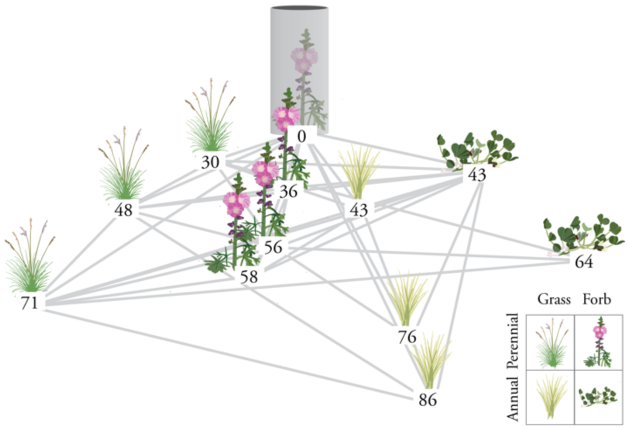


**Figure S2. Example of how networks were constructed for each plot.** The grey cylinder indicates the donor plant for the plot. Numbers beneath the receivers are the distance (in centimeters) from the donor. Degrees were calculated as how many plants each individual plant was connected to by shared fungal ASVs; for example, the perennial grass at 71 cm has 7 degrees. We visualized individual plot networks in CytoScape.

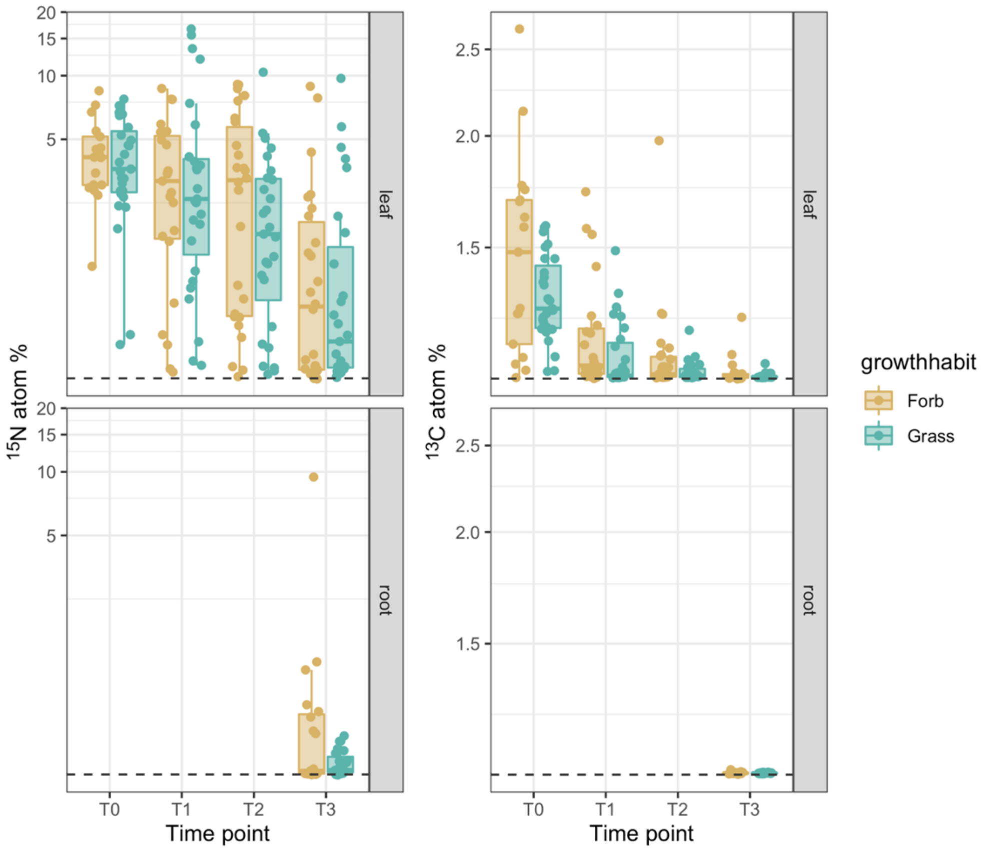


**Figure S3. Enrichment decreases over time in labelled plants.** Time points roughly correspond with T0 = time of labelling, T1 = 4 days later, T2 = 10 days later, T3 = 21 days later. Y-axis is in log10 scale.

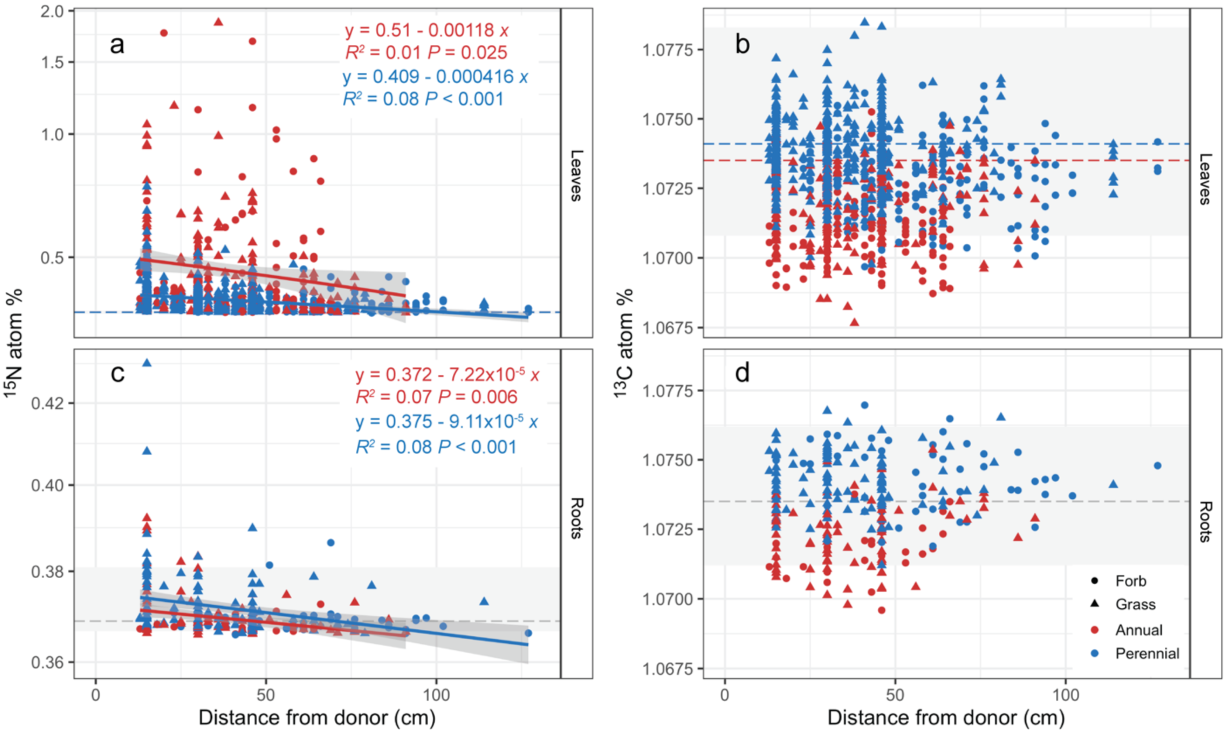


**Figure S4. Decreased receiver enrichment with distance from donor in leaves and roots.** Dashed lines indicate natural abundance means; grey boxes indicate range of natural abundance variation shown in Fig. 2. There is no systematic enrichment of $^{13}$C; however, there is high $^{15}$N enrichment in leaves. Y-axes are $\log_{10}$ scale. Note that the y-axis scales are different between $^{15}$N leaves and $^{15}$N roots. See Fig. 2 for boxplots of enrichment data by grass/forb and annual/perennial.

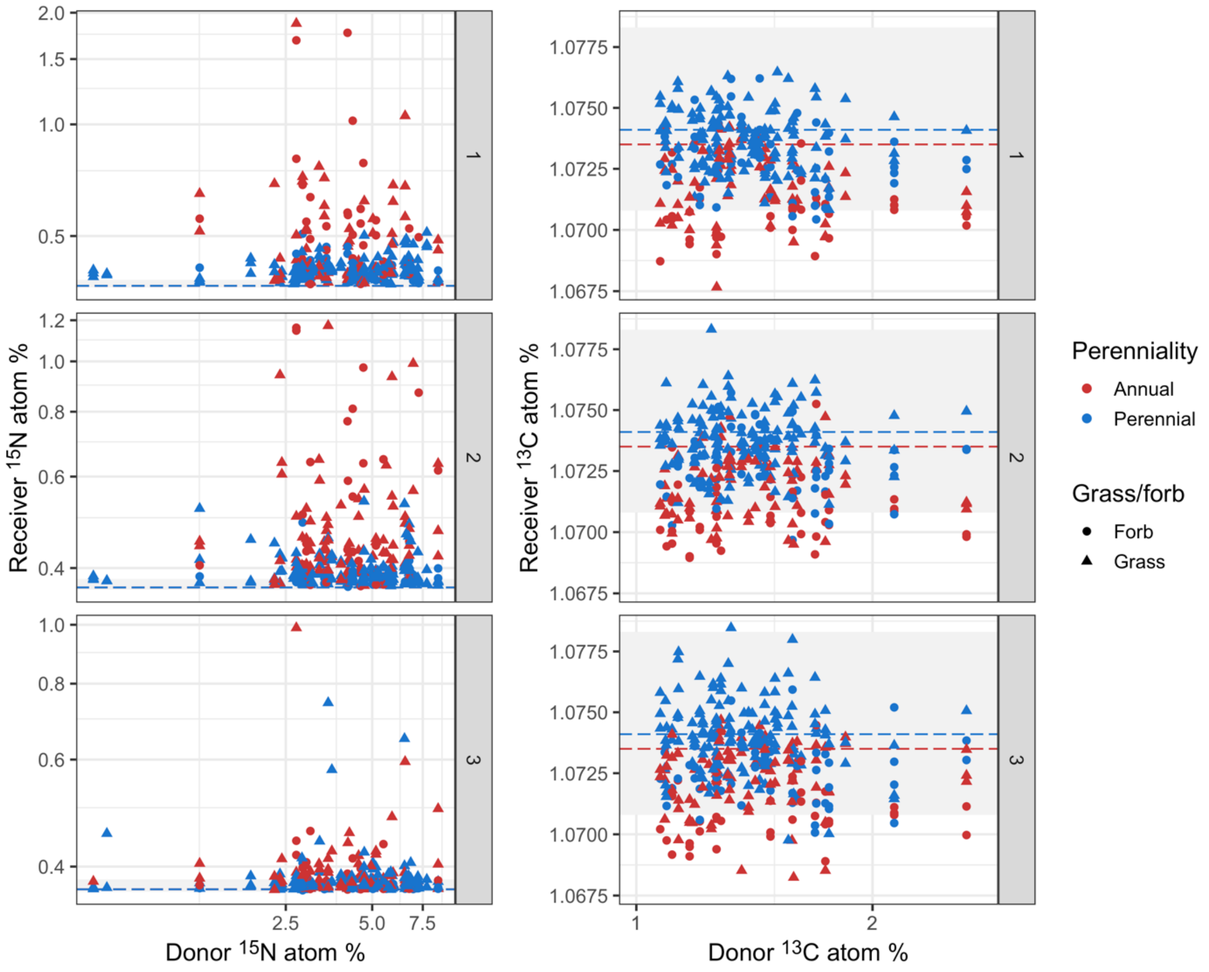


**Figure S5. Receiver enrichment did not correlate with initial donor enrichment.** Dashed lines indicate natural abundance means; grey boxes indicate range of natural abundance variation shown in Figure S1. (Grey boxes are too small to be noticeable for $^{15}$N.) $^{15}$N axes are log10 scaled. Note there is a sizeable difference between X- and Y-axes of both elements. Time point facets are time point 1 (~4 days post label), time point 2 (~10 days post label), and time point 3 (~21 days post label). Donors are all sampled at time point 0 (immediately after labelling).

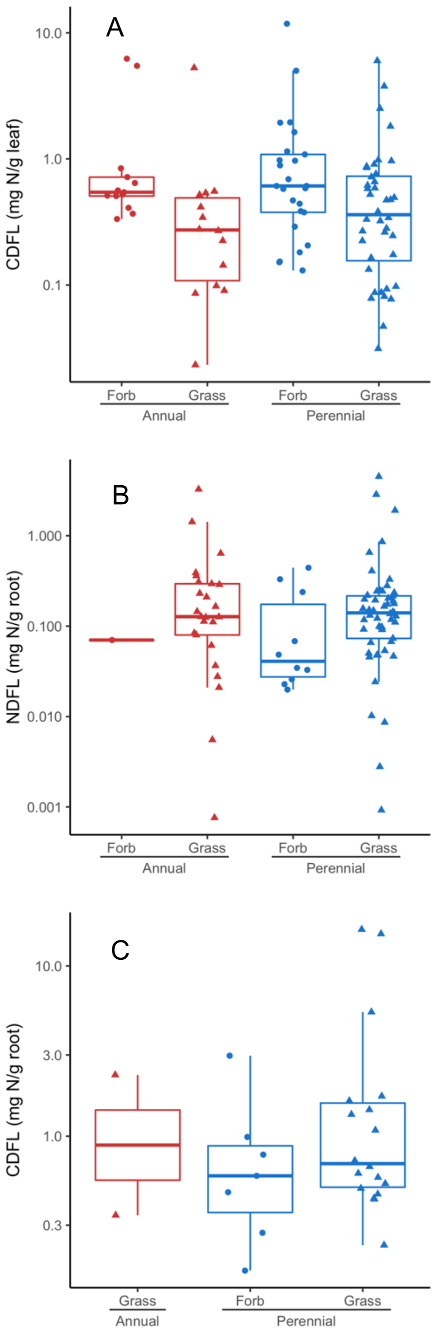


**Figure S6: Enrichment derived from label for A) CDFL in leaves, B) NDFL in roots, and C) CDFL in roots.** Points represent enriched individual receiver plants with associated DNA data at one time point of sampling. In A all three post-enrichment time points are shown; in B and C only time point 3 is shown. Y-axis is log10 scale. See Fig. S** for values over distance and Fig. 4 for NDFL in leaves.

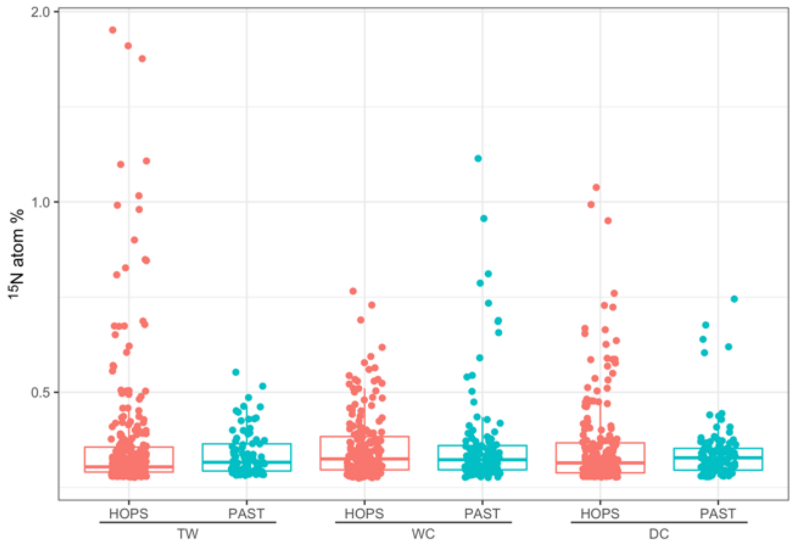


**Figure S7. No trend in $^{15}N$ enrichment across site and diversity treatment.** Y-axis is in log10 scale. HOPS indicates restored prairie sites, PAST indicates pasture sites. Sites are ordered from north (TW) to central (WC) to south (DC).

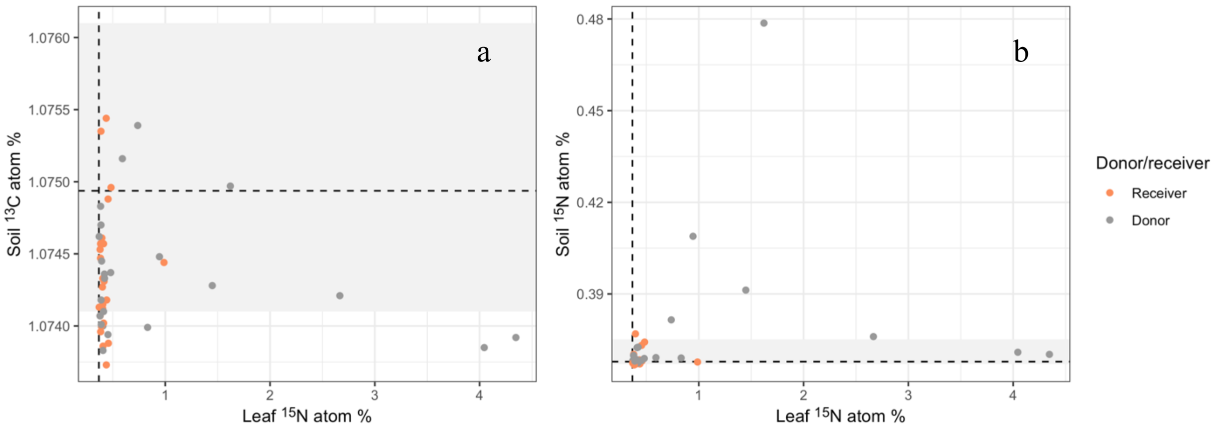


**Figure S8. Limited soil enrichment 21 days after labelling.** Dashed lines indicate average baseline enrichment. Grey boxes indicate range of baseline enrichment. Samples are subset to individuals most highly enriched with foliar $^{15}N$ at each site by restored prairie/introduced pasture as well as their associated donors.

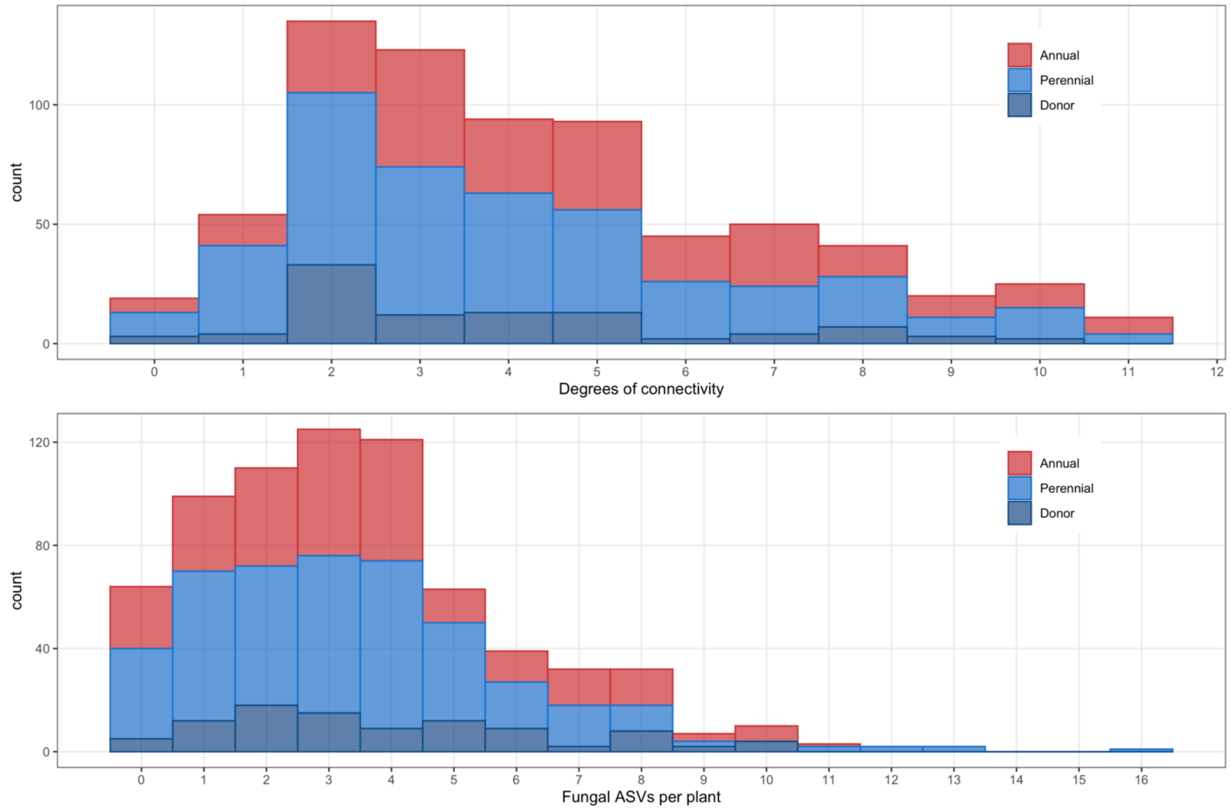


**Figure S9. Histogram of connectivity metrics showing left skew.** Degrees of connectivity indicate how many plants an individual shared at least one fungal ASV with in the same plot. Data are from time point 1 (approximately four days after labeling). Donors are the plants that we applied the isotopic label to.
